# Alternative polyadenylation characterizes epithelial and fibroblast phenotypic heterogeneity in pancreatic ductal adenocarcinoma

**DOI:** 10.1101/2021.09.18.460907

**Authors:** Swati Venkat, Michael E. Feigin

## Abstract

Human tumors are characterized by extensive intratumoral transcriptional variability within the cancer cell and stromal compartments. This variation drives phenotypic heterogeneity, producing cell states with differential pro- and anti-tumorigenic properties. While bulk RNA sequencing cannot achieve cell type specific transcriptional granularity, single cell sequencing has permitted an unprecedented view of these cell states. Despite this knowledge, we lack an understanding of the mechanistic drivers of this transcriptional and phenotypic heterogeneity. 3’ untranslated region alternative polyadenylation (3’ UTR-APA) drives gene expression alterations through regulation of 3’ UTR length. These 3’ UTR alterations modulate mRNA stability, protein expression and protein localization, resulting in cellular phenotypes including differentiation, cell proliferation, and migration. Therefore, we sought to determine whether 3’ UTR-APA events could characterize phenotypic heterogeneity of tumor cell states. Here we analyze the largest single cell human pancreatic ductal adenocarcinoma (PDAC) dataset and resolve 3’ UTR-APA patterns across PDAC cell states. We find that increased proximal 3’ UTR-APA is associated with PDAC progression and characterizes a metastatic ductal epithelial subpopulation and an inflammatory fibroblast population. Furthermore, we find significant 3’ UTR shortening events in cell state-specific marker genes associated with increased expression. Therefore, we propose that 3’ UTR-APA drives phenotypic heterogeneity in cancer.

## Background

Pancreatic ductal adenocarcinoma (PDAC) is a lethal disease with a 5-year survival rate of 10% [1]. PDAC tumors are characterized by a dense stroma and a high degree of cell type specific phenotypic variation that is integral to disease progression and drug resistance [2–4]. Over the past decade, bulk and single cell RNA sequencing (scRNA-seq) analyses uncovered substantial inter- and intratumoral transcriptional heterogeneity [5–7]. These studies have formed the basis for patient stratification and delineation of phenotypically distinct epithelial and stromal subpopulations. For example, tumor epithelial cells have been found to exist in subpopulations that exhibit differing proliferative and metastatic potential [6,8,9]. Similarly, phenotypically distinct subsets of cancer associated fibroblasts (CAFs) characterized by unique transcriptional profiles have been identified within the tumor microenvironment [10,11]. Two major CAF subclasses, inflammatory CAFs (iCAFs) and myofibroblastic CAFs (myCAFs), have distinct but crucial roles in tumor progression and therapeutic resistance [12,13]. However, mechanistic drivers of such transcriptional and phenotypic heterogeneity in PDAC remain unclear. Recently, we performed an in-depth analysis of sequencing data on PDAC tumors that established 3’ UTR alternative polyadenylation (APA) as a mechanistic driver of oncogene expression [14–16]. Specific PDAC oncogenes were found to undergo proximal 3’ UTR-APA (usage of proximal 3’ UTR polyadenylation site) resulting in shorter 3’ UTRs, driving increased expression. However, as this analysis made use of bulk RNA-seq data, it was impossible to determine the contribution of APA to cell type specific transcriptional heterogeneity. To determine if APA could be a mechanistic driver of phenotypic variation in cancer we now leverage the largest scRNA-seq human PDAC dataset recently published by Peng *et al*. [17]. Unlike bulk sequencing data, the majority of single cell sequencing protocols are 3’ biased, allowing robust detection of 3’-UTR-APA changes and the associated transcriptional heterogeneity in a high-resolution dataset [7,18–20]. To our knowledge, this is the first investigation of APA events associated with intratumoral heterogeneity.

## Results and Discussion

To understand whether APA is associated with cell type specific phenotypic variation, we sought to identify cell types that exhibit substantial 3’ UTR-APA events. To achieve this, we reanalyzed the scRNA-seq PDAC dataset (Additional file Fig. S1a) comprised of 11 normal pancreata and 24 tumor samples. We focused on cell types that form a significant proportion of the tumor, including acinar and ductal epithelial cells, and stromal fibroblasts and stellate cells. After quality control (see Methods), we processed a total of 22053 tumor cells across 21 tumor samples and 10345 normal cells across 11 pancreata for downstream analyses. We adapted a recently published algorithm to detect 3’ UTR-APA events from scRNA-seq data [18] (Additional file Fig. S1b). In concordance with previous findings, tumor tissues exhibited significantly higher proximal 3’ UTR-APA gene events (3’ UTR shortening) as compared to normal tissues [14,16]. In particular, tumor ductal cells showed significantly higher numbers of proximal 3’ UTR-APA events (1177 genes expressed shorter 3’ UTRs and 250 genes expressed longer 3’ UTRs) compared to normal ductal cells (Fig. 1a). While fibroblasts, acinar cells and stellate cells in PDAC tumors exhibited a higher number of proximal 3’ UTR-APA events compared to their normal counterparts, PDAC ductal cells exhibited the highest ratio of proximal to distal 3’ UTR-APA events (∼5:1) compared to other cell types. While a bulk PDAC RNA-seq study would reveal significant 3’ UTR-APA events occurring across a mixture of these cell types, it would fail to resolve cell type specific 3’ UTR-APA events. The extent of proximal 3’ UTR-APA in PDAC ductal cells motivated us to probe APA events within this transcriptionally diverse cell population. Peng and colleagues identified two subsets of PDAC ductal cells, namely ductal cell type 1 and ductal cell type 2. Ductal cell type 2 constituted the majority of the PDAC ductal cells and exhibited a malignant gene expression profile. Ductal cell type 1 expressed an abnormal gene expression profile that was distinct from the normal cells, representing a transcriptional state between normal and tumor ductal cells [17]. We performed dimensionality reduction and clustering to delineate these transcriptionally distinct subsets of normal and tumor ductal cells (Fig. 1b). Clustering revealed 6 transcriptionally distinct subclusters: normal ductal cells (dA), tumor ductal cell type 1 (dB) and tumor ductal cell type 2 (composed of subclusters dC, dD, dE, dF). Interpatient as well as intrapatient heterogeneity was detected in ductal cell type 2 with the majority of the patients represented in subcluster dC and a minority in subclusters dD, dE and dF (Additional file, Fig. S2a). Subcluster dE specific genes were enriched for metastatic markers (*HMGA1, ENO1, GABRP, IGFBP2, SDC1, LGALS1*) (Additional file, Fig. S2b) and pathway enrichment analysis of dE overexpressed genes showed epithelial to mesenchymal transition (EMT) as a top hit supporting its metastatic phenotype (Additional file, Fig. S2c) [21–26]. In contrast, gene expression and pathway analysis of subcluster dD specific genes showed enrichment for well-differentiated PDAC markers (*REG4, TFF1, TFF2, TFF3, VSIG2, LGALS4*), highlighting the extensive phenotypic heterogeneity exhibited by PDAC ductal cells (Additional file, Fig. S2d) [6].

**Figure 1.**
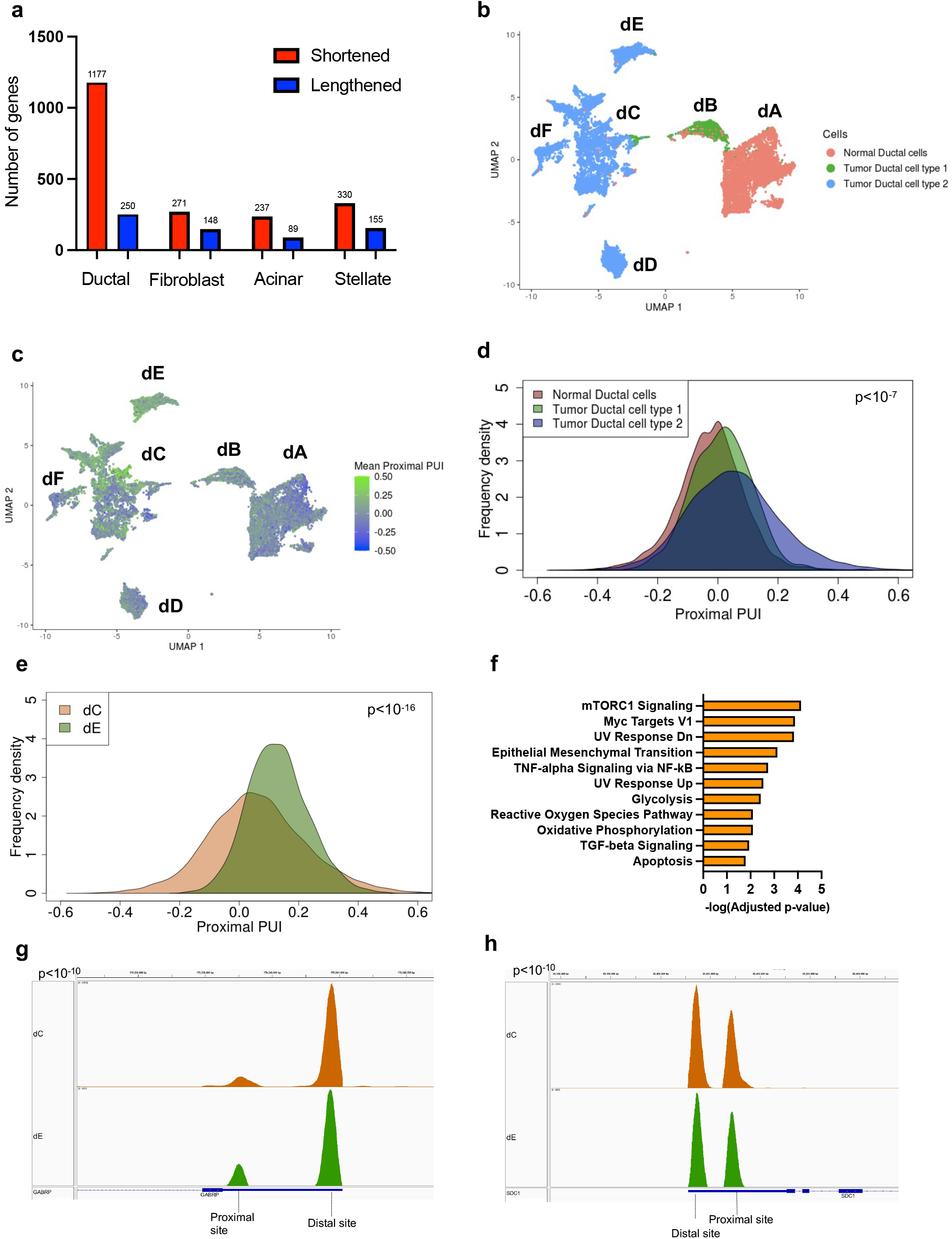
Proximal APA in tumor epithelium is associated with PDAC progression and malignant phenotypes. 1a. A plot of number of shortened (red) and lengthened (blue) 3’ UTR-APA events across four PDAC cell types compared to their counterparts in normal pancreas. 1b. UMAP embedding of ductal cells (dots) from normal pancreata and tumor patients. Color indicates the ductal cell type membership. Notations dA-dF denote the subclusters. 1c. UMAP embedding of ductal cells from normal pancreata and tumor patients. Color indicates degree of mean proximal PUI in each cell (blue, low; green, high). 1d. Distribution of mean proximal PUI of single cells in normal ductal cells (orange), tumor ductal cell type 1 (green) and ductal cell type 2 (blue) (every pairwise comparison yielded p<10^−7^). 1e. Distribution of mean proximal PUI of single cells in subcluster dE (green) compared to subcluster dC (brown) (p<10^−16^). 1f. Significantly enriched pathways (FDR < 0.01) associated with 3’ UTR altered genes between subclusters dE and dC. 1g. IGV plot highlighting the 3’ UTR density profile differences of the metastatic gene *GABRP* between subclusters dC (brown) and dE (green). 1h. IGV plot highlighting the 3’ UTR density profile differences of the metastatic gene *SDC1* between subclusters dC (brown) and dE (green).

We first sought to characterize APA patterns across the broad ductal cell states (normal ductal cells, tumor ductal cell type 1, tumor ductal cell type 2) to determine the relationship between APA and tumor progression. We determined the mean proximal polyA site usage index (mean proximal PUI), the extent of 3’ UTR proximal site usage for each cell, averaged over all genes (see Methods, [18]). A higher mean proximal PUI indicates enhanced cleavage at proximal polyadenylation sites in the cell (resulting in shorter 3’ UTRs). We plotted the mean proximal PUI for every ductal cell associated with each cell state (Fig. 1c). Pseudotime analysis confirmed progression from a normal state (normal ductal cells) to an abnormal intermediate state (tumor ductal type 1) to a malignant ductal state (tumor ductal type 2) (Additional file, Fig. S2e). This malignant progression was associated with a progressive and significant increase in mean proximal PUI (Fig 1d). Therefore increased proximal 3’-UTR-APA is associated with malignant progression in PDAC.

We noted substantial APA heterogeneity within the subclusters comprising tumor ductal cell type 2 (dC, dD, dE) and therefore quantified proximal 3’ UTR-APA patterns across these cells. The cells in the metastatic subcluster dE showed a significant increase in mean proximal PUI compared to dC, indicating increased 3’ UTR shortening events in dE (Fig. 1e). In contrast, the cells in the well-differentiated PDAC subcluster dD showed a significant decrease in mean proximal PUI compared to dC, indicating decreased 3’ UTR shortening events (Additional file, Fig. S2f). To determine if these APA events are associated with known metastatic driver genes, we performed pathway enrichment analysis of the 3’ UTR altered genes in dE, which revealed EMT as a top hit (Fig. 1f). Furthermore, we found significantly increased proximal APA of metastasis-promoting genes preferentially expressed in dE, including *GABRP* and *SDC1* (Fig. 1g, 1h, Additional file, Fig. S2b). This suggests a novel role of proximal 3’ UTR-APA in orchestrating the metastatic PDAC phenotype.

CAFs are a transcriptionally and phenotypically heterogeneous population in the tumor microenvironment that make fundamental contributions to both progression and therapy response [11,12,27–29]. How this transcriptional heterogeneity is developed and maintained during tumorigenesis is integral to the advancement of more effective therapeutic strategies. In PDAC, two major CAF subtypes have been discovered and functionally characterized – myCAFs, responsible for secreting the extracellular matrix components that promote a dense desmoplastic stroma, and iCAFs, responsible for secreting IL-6 and other inflammatory mediators. To investigate the role of APA in CAF biology, we clustered normal fibroblasts and CAFs and identified transcriptionally differing subclusters within the CAF population (Fig. 2a). Clustering revealed transcriptionally distinct subclusters including normal fibroblast cells (fA) and tumor fibroblast cells (composed of subclusters fB-fE). Pathway analysis and cluster specific gene markers revealed fC as a myCAF population (*ACTA2, POSTN, MMP11, IGFBP3, COL12A1, THBS2*), (Additional file, Fig. S3a, S3c) and fD as an iCAF population (*HAS1, HAS2, CCL2, UGDH, SOD2, LMNA*), (Additional file, Fig. S3b, S3d) [12,13]. To characterize 3’ UTR-APA patterns, we determined the mean proximal PUI for every normal and tumor fibroblast cell (Fig. 2b). In contrast to normal fibroblasts, the tumor fibroblast population showed a small but significant increase in proximal 3’ UTR-APA (Additional file, Fig. S3e), indicative of more 3’ UTR shortening events in CAFs. We next examined 3’ UTR-APA underlying CAF heterogeneity and found no significant difference between normal fibroblasts and the myCAF population (Fig. 2c). In contrast, there was a significant increase in 3’ UTR shortening in the iCAF population (Fig. 2d, Additional file, Fig. S3f), revealing that increased proximal APA characterizes the inflammatory CAF phenotype. Importantly, we found significant increased proximal APA of critical iCAF markers such as *SOD2* and *UGDH* associated with their increased expression in iCAFs (Fig. 2e, 2f, Additional file, Fig. S3b). This suggests a novel role of 3’ UTR-APA in orchestrating the inflammatory CAF phenotype.

**Figure 2.**
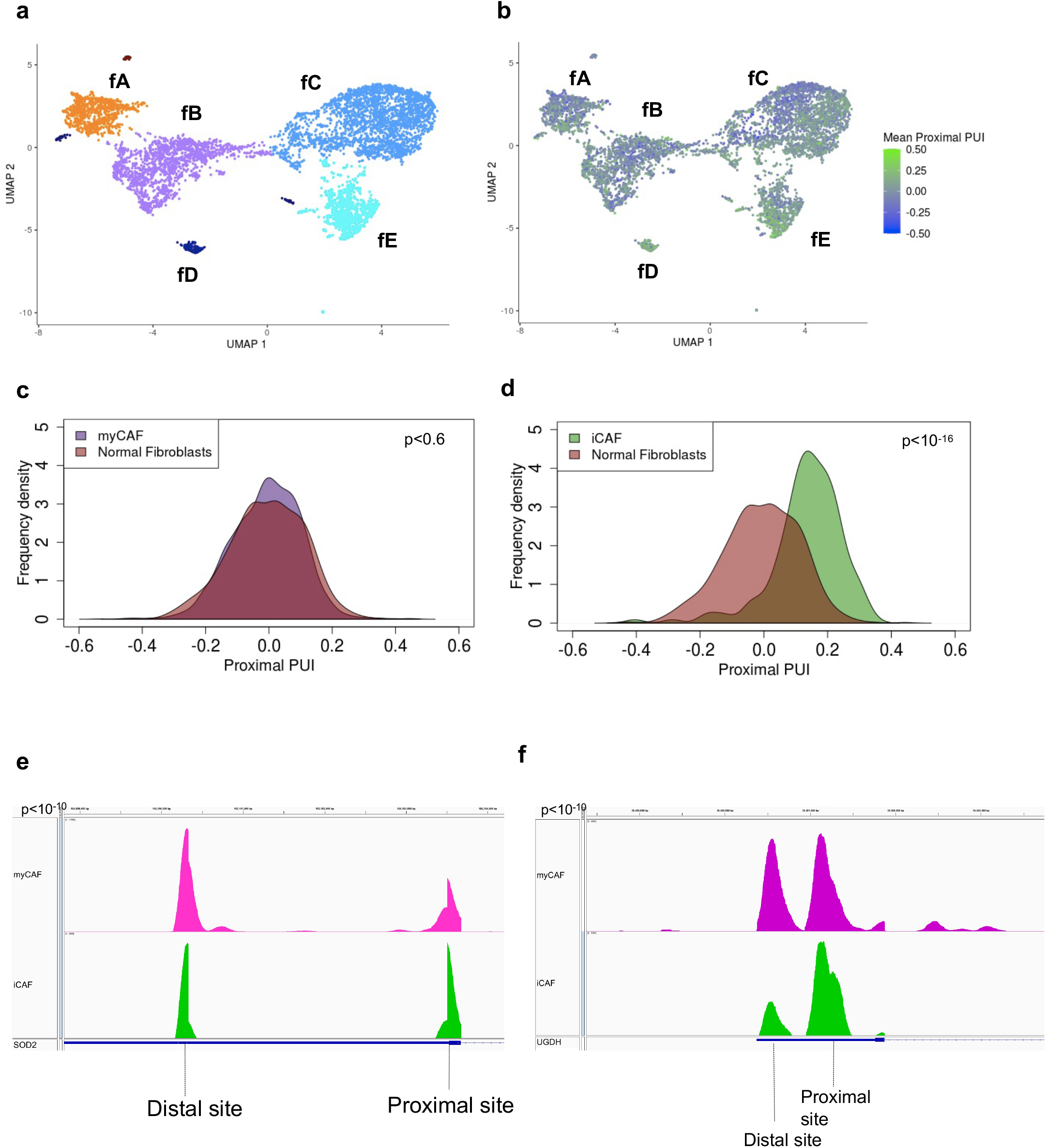
Increased proximal APA characterizes the inflammatory CAF phenotype. 1a. UMAP embedding of fibroblast cells (dots) from normal pancreata and tumor patients. Color indicates the fibroblast cell type membership. Notations fA-fE denote the subclusters. 1b. UMAP embedding of fibroblast cells from normal pancreata and tumor patients. Color indicates degree of mean proximal PUI in each cell (blue, low; green, high). 1c. Distribution of mean proximal PUI of single cells (p=0.6) in normal fibroblast cells (orange) and myCAFs (purple). 1d. Distribution of mean proximal PUI of single cells (p<10^−16^) in iCAFs (green) compared to normal fibroblast cells (orange). 1e. IGV plot highlighting the 3’-UTR density profile differences of the iCAF activated transcription factor *SOD2* between iCAFs (green) and myCAFs (purple). 1f. IGV plot highlighting the 3’-UTR density profile differences of the iCAF marker *UGDH* between iCAFs (green) and myCAFs (purple).

## Conclusions

3’ UTR-APA is an underappreciated driver of gene dysregulation in cancer. Single cell sequencing has revealed that tumors have high degrees of transcriptional and phenotypic heterogeneity, both within the cancer cell and stromal compartments. However, drivers of such complex phenotypic heterogeneity remain unclear. In this study, we investigated 3’UTR-APA associated phenotypic heterogeneity using single cell data. To our knowledge, this is the first investigation of APA events associated with intratumoral heterogeneity. We demonstrate that 3’ UTR shortening increases progressively during PDAC progression. Furthermore, 3’ UTR shortening of critical metastatic and iCAF marker genes is associated with increased expression, thereby defining cell identity. Increased proximal 3’ UTR-APA characterizes a metastatic ductal subpopulation in tumor epithelial cells as well as an inflammatory CAF population in the PDAC stroma. We propose that 3’ UTR-APA drives phenotypic heterogeneity both in the tumor epithelium and within the tumor microenvironment.

## Methods

### Bioinformatic processing of human scRNA-seq data

scRNA-seq FASTQ files of 24 PDAC patients and 11 normal pancreata were downloaded from Genome Sequence Archive (GSA) (Accession: CRA001160, Bioproject: PRJCA001063). Cell Ranger 3.1.0 using standard parameters was used to align each file to the hg19 genome [30]. Appropriate chemistry and alignment by Cell Ranger was detected for 21 patients and 11 normal tissues and these data were used for downstream analyses. We focused on annotated cells (Peng *et al*. [17]) with at most 6000 genes/cell (to eliminate doublets) and with at least 200 genes/cells. Cells with >10% mitochondrial counts and genes occurring in <3 cells were excluded from the analysis. This yielded 10345 normal cells and 22053 tumor cells for the analysis of 3’ UTR-APA events.

### Analysis of 3’UTR-APA events

Analysis of 3’ UTR-APA events was performed by manual implementation of the scRNA-seq algorithm proposed in [18] (Fig. S1b). Briefly, PCR duplicates were discarded from aligned BAM files using UMI tools [31]. These files were used to detect peaks in 3’ UTR read density using Homer findPeaks function [32,33]. Additionally, cell type identity was obtained from Peng *et al*. [17] and we used this information to annotate major cell types and generate cell type specific BAM files. Reads in cell type specific BAMs that mapped to Homer-determined peak positions were measured using Feature counts (Rsubread package) [34]. Low count peaks (<10 CPMs over all cell clusters) and peaks with A-rich sequences [18] were filtered out, allowing identification of statistically significant 3’UTR-APA events and mean proximal PUI at a single cell level exactly as described in [18]. IGV plots were used to visualize the read density changes for the 3’ UTR altered genes. Frequency density plots were used to visualize distribution of mean proximal PUI across single cells in a subcluster and significant differences between subclusters were assessed using the Wilcoxon ranked sum test with continuity correction.

### Bioinformatics analyses and statistical methods

Subsequent analyses were carried out in R 4.0.4. Monocle3 was used to analyze single cell trajectories to determine cell state transitions [35]. Top 200 differentially expressed genes between normal and tumor type 2 ductal cells were used for dimensionality reduction via UMAP and clustering and the mean proximal PUI for each cell was overlayed. The top 25 cluster-specific marker genes were identified using the top_markers function in Monocle3. Differentially expressed genes between the subclusters were identified using the FindMarkers function in Seurat4 [36]. Gene Set Enrichment Analysis (GSEA) and Enrichr were used to perform pathway analysis using the MSigDB hallmark, KEGG and Reactome gene sets [37,38]. Enrichment of the input genes (3’-UTR-APA altered genes/differentially expressed genes) in Enrichr was computed using the Fisher’s exact test and p-values were adjusted using the Benjamini-Hochberg correction (FDR < 0.01). A similar approach was implemented for analysis of fibroblasts.

### Availability of data and materials

The R code written for this analysis is available on GitHub (https://github.com/feiginlab/APA_PDA).

## Acknowledgments

This work was supported by NCI grant P30 CA016056, and an award from the Roswell Park Alliance Foundation to MEF. We thank the members of the Feigin laboratory and Dr. Ethan Abel for their helpful comments on the manuscript.

## Figure legends

**Figure S1.**
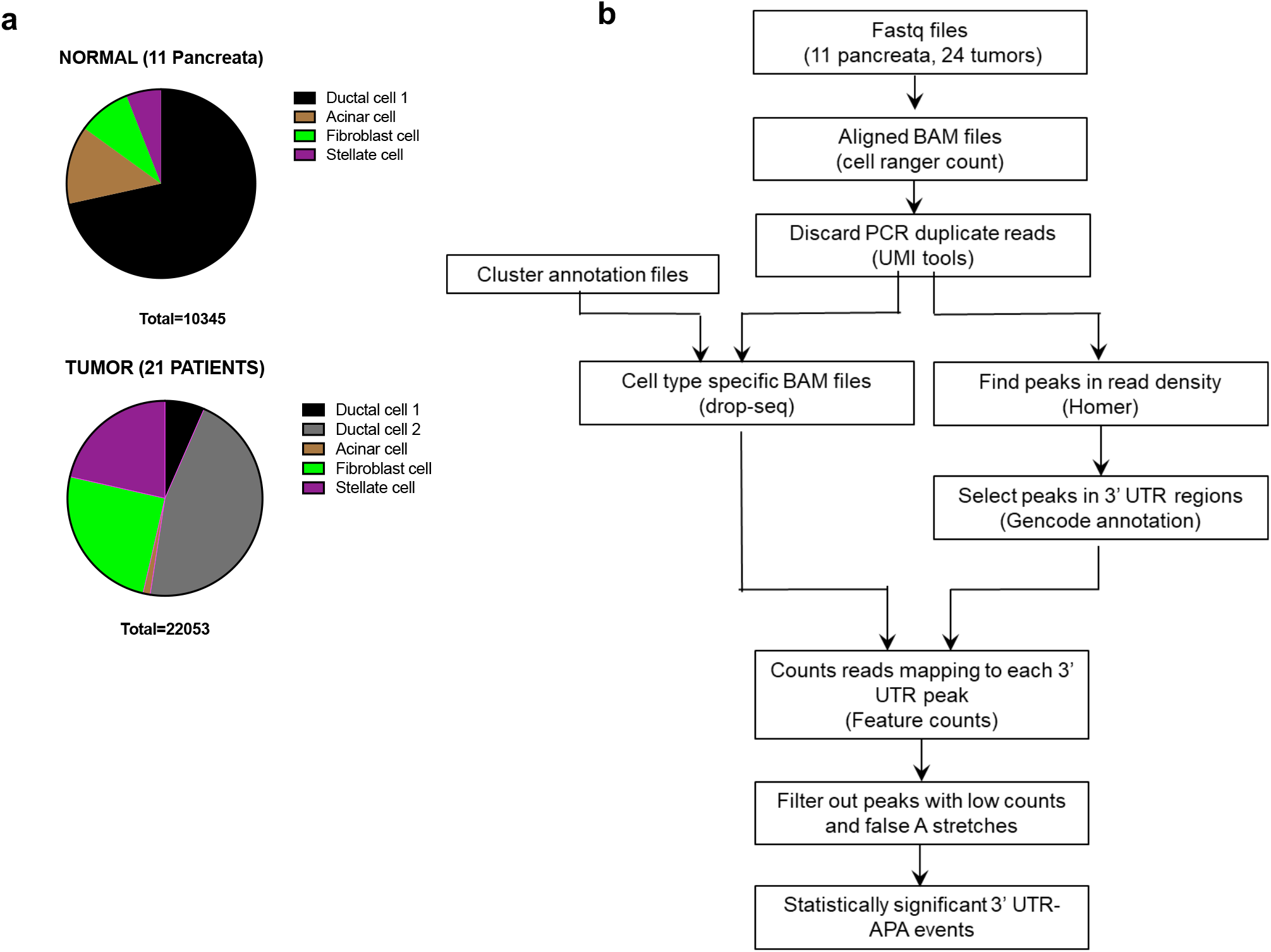
Description of the scRNA-seq dataset and the workflow used to quantify 3’ UTR-APA in PDAC. S1a. A pie graph representing the single cell dataset that was used for downstream analyses of 3’ UTR-APA patterns. Proportion of ductal and acinar cells in the epithelium, and fibroblast and stellate cells in the stroma are highlighted. S1b. The workflow implemented to detect and quantify 3’ UTR-APA events from single cell sequencing data (adapted from [18]).

**Figure S2.**
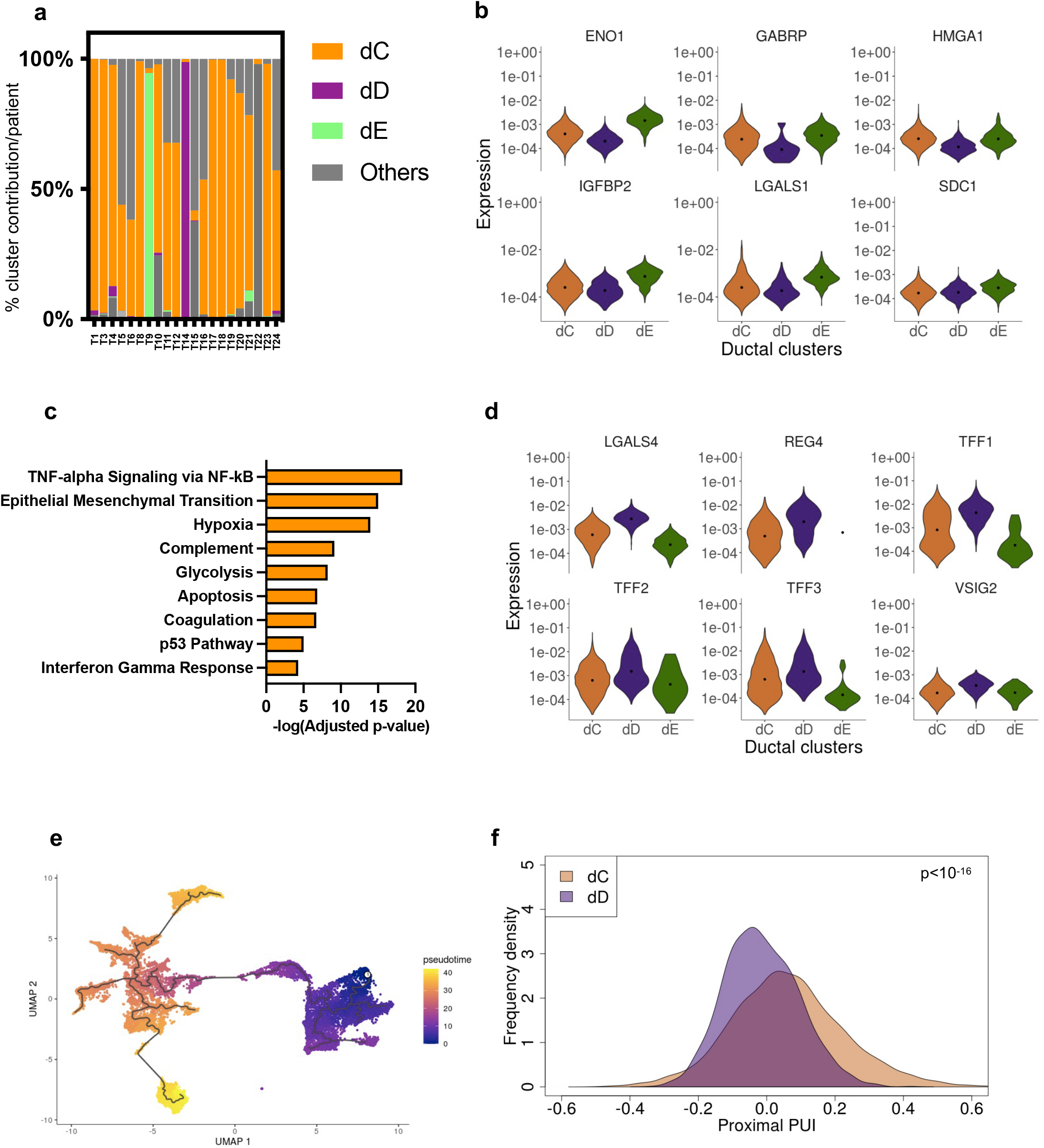
Proximal APA in tumor epithelium is associated with PDAC progression and malignant phenotypes. S2a. Barplot showing contribution of different ductal subclusters to each PDAC patient. S2b. Violin plots of select metastatic markers across ductal cell type 2 subclusters (p<0.001). S2c. Significant enriched pathways (FDR < 0.01) associated with genes overexpressed in dE compared to dC. S2d. Violin plots of select well-differentiated PDAC markers across ductal cell type 2 subclusters (p<0.001). S2e. Pseudo-time analysis depicting progression of ductal cell states (purple, early; yellow, late) based on their gene expression profiles. S2f. Distribution of mean proximal PUI of single cells in subcluster dD (purple) compared to subcluster dC (brown) (p<10^−16^).

**Figure S3.**
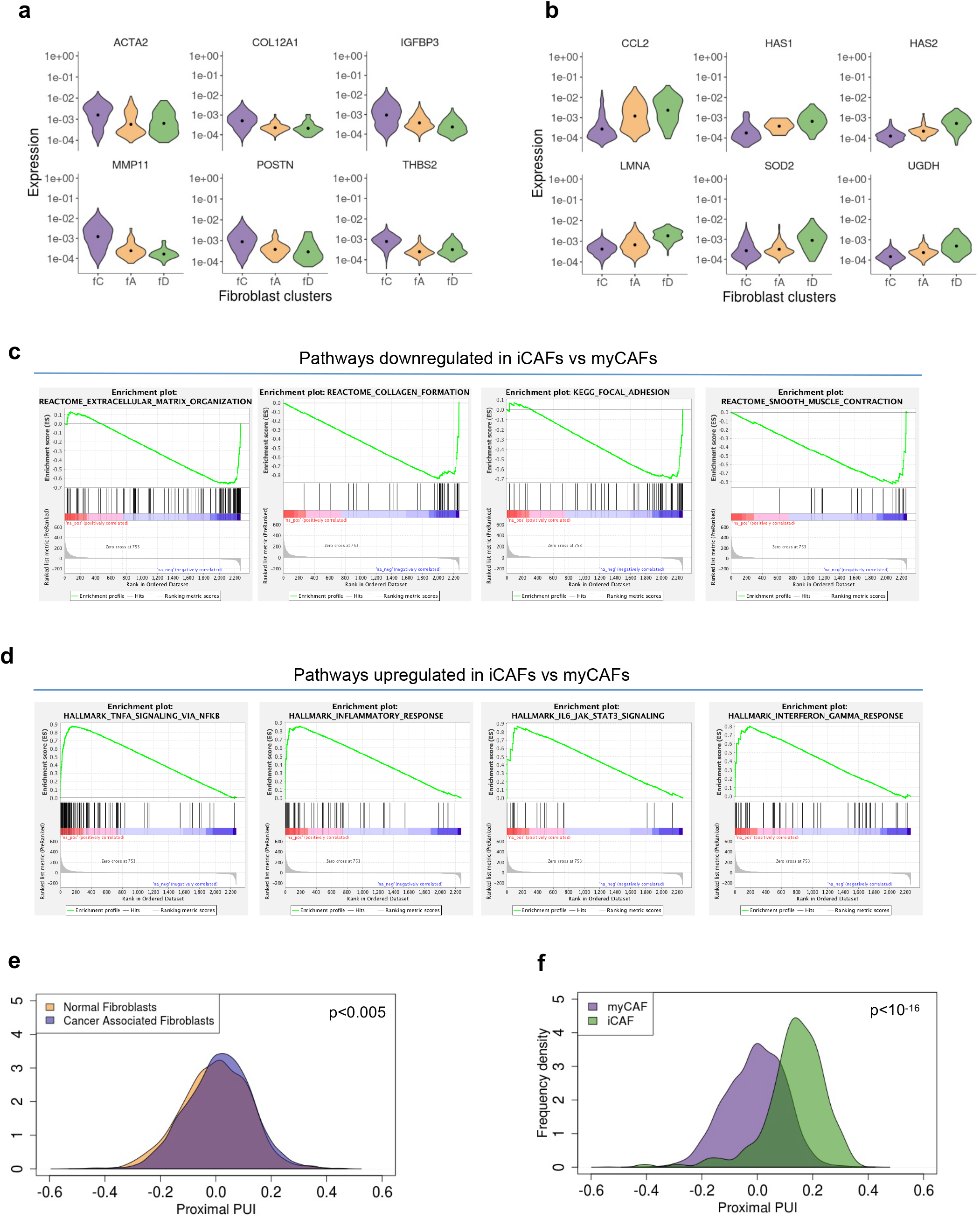
Increased proximal APA characterizes the inflammatory CAF phenotype. S3a. Violin plots of myCAF markers across normal fibroblasts (fA, orange) and specific tumor fibroblast subclusters (fC, purple; fD, green). S3b. Violin plots of iCAF markers across normal fibroblast (fA) and specific tumor fibroblast subclusters (fC, fD). S3c. GSEA of significantly downregulated pathways in iCAFs compared to my CAFs. S3d. GSEA of significantly upregulated pathways in iCAFs compared to my CAFs. S3e. Distribution of mean proximal PUI of single cells of normal fibroblasts (orange) compared to tumor fibroblasts (blue) (p<0.005). S3f. Distribution of mean proximal PUI of single cells of iCAFs (green) compared to myCAFs (purple) (p<10^−16^).

